# A bacterial effector uncovers a metabolic pathway involved in resistance to bacterial wilt disease

**DOI:** 10.1101/2020.09.22.307595

**Authors:** Yaru Wang, Rafael J. L. Morcillo, Gang Yu, Achen Zhao, Hao Xue, Jose S. Rufian, Yuying Sang, Alberto P. Macho

## Abstract

Bacterial wilt caused by the soil-borne pathogen *Ralstonia solanacearum* is a devastating disease worldwide. Upon plant colonization, *R. solanacearum* replicates massively, causing plant wilting and death; collapsed infected tissues then serve as a source of inoculum. In this work, we show that the metabolic pathway mediated by pyruvate decarboxylases (PDCs), activated in response to low oxygen and involved in drought stress tolerance, contributes to resistance against bacterial wilt disease. Arabidopsis and tomato plants with deficient PDC activity are more susceptible to bacterial wilt, and treatment with either pyruvic acid or acetic acid (substrate and product of the PDC pathway, respectively) enhances resistance. An effector protein secreted by *R. solanacearum*, RipAK, interacts with PDCs and inhibits their oligomerisation and enzymatic activity. This work reveals a metabolic pathway involved in resistance to biotic and abiotic stresses, and a bacterial virulence strategy to promote disease and the completion of the pathogenic life cycle.

## Introduction

Environmental stresses have a strong impact on plant development and survival, and are therefore a serious threat to crop production. To cope with stress, plant cells are equipped with a sophisticated network of receptors, signalling pathways, and physiological responses that allow the integration of multiple and often simultaneous environmental signals to adapt to their changing environment. Although our understanding of the plant signalling pathways associated to stress (both biotic and abiotic) and metabolic adaptations has significantly expanded over the past few years, the association between these pathways is still poorly understood, and often limited by their man-made classification as responsive to one or another type of stress.

The bacterial plant pathogen *Ralstonia solanacearum* is the causal agent of the bacterial wilt disease in more than 250 plant species, including economically important crops, such as tomato, potato, pepper, eggplant, or banana (Elphinstone et al., 2005; Mansfield et al., 2012). As a soil-borne bacterium, *R. solanacearum* enters plants through the roots, invades the xylem vessels, and rapidly colonizes the whole plant (Xue et al, 2020). *R. solanacearum* shows a hemi-biotrophic behaviour, proliferating in live tissues in early stages of the infection; subsequently, massive bacterial replication and the production of large amounts of exopolysaccharide lead to clogging of the xylem vessels and vascular dysfunction, eventually causing plant wilting and death (Genin, 2010; Mansfield et al., 2012). Before its death, an infected plant can host a huge bacterial population, reaching up to 10^10^ colony-forming units (cfu) per gram of tissue (Genin, 2010). Therefore, the wilting and collapse of plant tissues bring back to the soil an extremely concentrated bacterial inoculum for additional potential host plants, thus perpetuating the pathogenic cycle of *R. solanacearum*.

The best-studied plant defence mechanisms against invading pathogens rely on the perception of microbial molecules that are considered as invasion patterns (Cook et al., 2015). Highly conserved and abundant microbial molecules, often involved in housekeeping microbial functions, can be perceived by plants as pathogen-associated molecular patterns (PAMPs), and are notorious elicitors of plant immune responses (Boller and Felix, 2009). *R. solanacearum* PAMPs identified to date include the elongation factor *Tu* and cold-shock proteins (Lacombe et al., 2010; Wei et al., 2018). Most gram-negative bacterial pathogens use a type-III secretion system (T3SS) to inject effector proteins (type-III effectors; T3Es) inside plant cells. T3Es exert virulence activities aimed at promoting bacterial proliferation, such as the suppression of immunity or the manipulation of other plant functions (Macho, 2016; Macho and Zipfel, 2015; Toruño et al., 2016). However, resistant plants harbouring specific intracellular receptors can detect specific T3Es or their activities, activating immune responses (Chiang and Coaker, 2015). The T3E repertoire of *R. solanacearum* is particularly diverse: a single strain can inject more than 70 different T3Es inside plant cells (Sabbagh et al., 2019). Given that microbial effectors have evolved to target plant cellular functions that are important during plant-microbe interactions, they can be used as probes to identify and characterize plant cellular functions that contribute to disease resistance or susceptibility (Toruño et al., 2016). One of the T3Es in the *R. solanacearum* repertoire, RipAK (also known as Rip23 (Mukaihara et al., 2010), is broadly conserved among strains from the phylotypes I and III (Sabbagh et al., 2019) (https://iant.toulouse.inra.fr/T3E), suggesting an important role in the pathogenicity of strains with the same phylogenetic origin. RipAK has been reported to localize at peroxisomes in protoplasts of *Arabidopsis thaliana* (hereafter, Arabidopsis), inhibiting host catalases to suppress plant immunity in tobacco (Sun et al., 2017). In this work, we found that RipAK localizes to the cytoplasm in *Nicotiana benthamiana* cells, in addition to forming speckles that partially overlap with peroxisomes. We show that, despite the redundancy expected among *R. solanacearum* T3Es, RipAK contributes significantly to the development of disease in Arabidopsis and tomato plants upon soil-drenching inoculation with *R. solanacearum*.

In plant cells, RipAK associates with plant pyruvate decarboxylases (PDCs), which are metabolic enzymes involved in fermentation under low oxygen conditions, and inhibits PDC enzymatic activity. Further genetic analysis showed that PDCs contribute to plant resistance against bacterial wilt, and chemical treatments showed that different organic acids in the PDC pathway, including pyruvate and acetate, enhance plant resistance against bacterial wilt. This work therefore reveals a novel pathway involved in disease resistance, which is inhibited by a *R. solanacearum* T3E, thus promoting the completion of the pathogenic life cycle.

## Results and discussion

### RipAK contributes to *R. solanacearum* infection in Arabidopsis and tomato

RipAK is highly conserved in *R. solanacearum* strains from the phylotypes I and III (recently named *R. pseudosolanacearum*; Sabbagh et al., 2019), which include the reference GMI1000 strain. Such conservation suggests an important role of RipAK in the pathogenicity of *R. pseudosolanacearum* strains. To determine the contribution of RipAK to bacterial wilt caused by GMI1000, we generated a *ΔripAK* knockout mutant (Figures S1A and S1B). Upon soil-drenching inoculation, the *ΔripAK* mutation reduced significantly the ability of *R. solanacearum* to cause disease symptoms in Arabidopsis (Figures 1A, 1B, S1C, and S1D) and tomato (Figures 1C, 1D, S1E, and S1F), which is a natural and agronomically important host for *R. solanacearum* (Hayward, 1991). In both cases, such virulence attenuation was rescued by the complementation of *ripAK* in the mutant background (Figures 1A-D and S1A-F), indicating that, despite the large number of T3Es secreted by *R. solanacearum*, RipAK plays a significant role in bacterial virulence. Interestingly, we did not detect attenuation in the replication of the *ΔripAK* mutant upon bacterial injection in the stem of tomato plants (Figure S1G). Although stem injection is a more aggressive inoculation method that bypasses the root penetration and colonization process, the same experimental setup has allowed us to detect significant growth attenuation of T3E mutants impaired in the generation of bacterial nutrients or the suppression of plant immunity (Xian et al., 2019; Yu et al., 2019). Therefore, these results may suggest that, rather than being required for bacterial multiplication, the RipAK virulence activity contributes to the development of disease symptoms or the initial penetration through the root.

**Figure 1.**
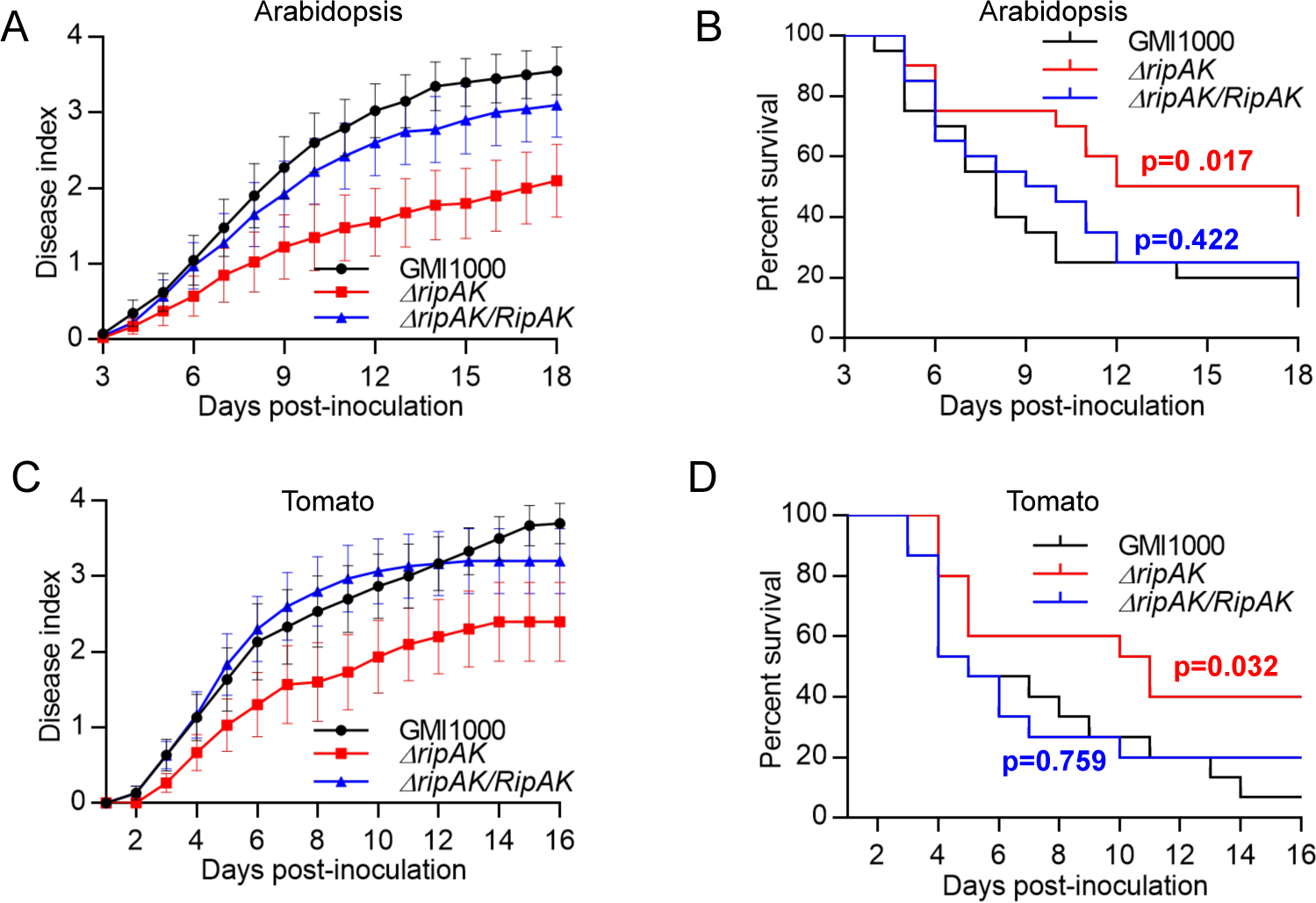
RipAK contributes to *R. solanacearum* infection. *R. solanacearum* soil-drenching inoculation assays in Arabidopsis (A, B) and tomato (C, D) performed with GMI1000 WT, *ΔripAK* mutant, and RipAK complementation (*ΔripAK/RipAK*) strains. n≥15 plants per genotype (for Arabidopsis) or n≥12 plants per genotype (for tomato). In A and C, the results are represented as disease progression, showing the average wilting symptoms in a scale from 0 to 4 (mean ± SEM). B and D show the survival analysis of the data in A and C, respectively; the disease scoring was transformed into binary data with the following criteria: a disease index lower than 2 was defined as ‘0’, while a disease index equal or higher than 2 was defined as ‘1’ for each specific time point. Statistical analysis was performed using a Log-rank (Mantel-Cox) test, and the corresponding p value is shown in the graph with the same colour as each curve. Nine and five independent biological replicates were performed for inoculations in Arabidopsis and tomato, respectively, and composite data representations are shown in Figure S1C-F.

### RipAK subcellular localization in plant cells

In order to understand the mode of action of RipAK in plant cells, we first studied the subcellular localization of a RipAK-GFP fusion protein expressed in *N. benthamiana* leaves using *Agrobacterium tumefaciens* (hereafter, *Agrobacterium*). RipAK-GFP localized in speckles and in the cytoplasm of plant cells (Figure S2A). Western blot analysis did not show a detectable amount of cleaved GFP in the experimental conditions used in these assays (Figure S2B), suggesting that the observed fluorescence corresponds to the RipAK-GFP fusion. The observed cytoplasmic localization is in contrast with the previous observation that RipAK-GFP localizes in peroxisomes when transiently expressed in Arabidopsis protoplasts (Sun et al., 2017), although such study employed a shorter RipAK version with an N-terminal truncation of 70 amino acids in comparison with the conserved RipAK reference sequence used in this work (Sun et al., 2017; Figures S2C and S2D). We used GFP immunoprecipitation (IP) followed by mass spectrometry (MS) analysis to verify that the full RipAK-GFP (including the aforementioned 70 amino acids at the N-terminal) indeed accumulated in plant cells in our assays (Figure S2E). RipAK-GFP co-expression with the peroxisome marker PTS1 (Goedhart et al., 2012) fused to an mTurquoise2 fluorescent tag (mT-PTS1) showed that only a subgroup of the RipAK-GFP speckles co-localized with the peroxisome marker, while others did not (Figure S2A). Then, in order to compare both RipAK versions, we generated a RipAK^Δ1-70^-GFP truncated version (RipAK^71-809^), equivalent to that used by Sun *et al* (2017), with a predicted molecular weight approximately 7 kDa smaller than the full RipAK (Figures S2B and S2C). Both wild-type (WT) and truncated versions showed similar localization in the cytoplasm and fluorescent speckles (Figures S2A and S2B), suggesting that the different results in our work and that by Sun *et al* are due to the different experimental systems used.

### RipAK interacts with pyruvate decarboxylases (PDCs)

To identify protein targets of RipAK in plant cells, we performed a yeast two-hybrid (Y2H) screen using RipAK as a bait against a library of cDNA from tomato roots inoculated with *R. solanacearum*, obtaining numerous colonies containing different fragments of a tomato gene encoding a homolog of Arabidopsis *PYRUVATE DECARBOXYLASE* (*PDC*) genes (Table S1, Figure S3A). Arabidopsis has three genes annotated as *PDCs*, which encode predicted cytoplasmic proteins, and AtPDC1-GFP was shown to localize in the cytoplasm in Arabidopsis (Rasheed et al., 2018); accordingly, RipAK-GFP co-localized with different RFP-tagged AtPDCs in the cytoplasm upon transient expression in *N. benthamiana* (Figure S3B). The interaction between RipAK and SlPDC2 (identified in the Y2H screen; Figure S3A) was confirmed *in planta* by coIP of RipAK tagged with hemagglutinin (HA) and SlPDC2-GFP transiently expressed in *N. benthamiana* (Figure 2A). The association between RipAK-HA and AtPDCs-GFP (AtPDC1, AtPDC2, and AtPDC3) was also detected by coIP (Figure 2B), and direct interaction between these proteins was confirmed by Split-Luciferase (Split-LUC) assays (Figures 2C and S3C). Intriguingly, upon IP of either SlPDC2-GFP or AtPDCs-GFP, we found an additional immunoprecipitated band of RipAK-HA, which was approximately 20-25 kDa smaller than the original RipAK-HA (Figures 2A and 2B). Since that smaller band was not present in crude extracts, it is possible that RipAK undergoes N-terminal cleavage upon interaction with PDCs in plant cells.

**Figure 2.**
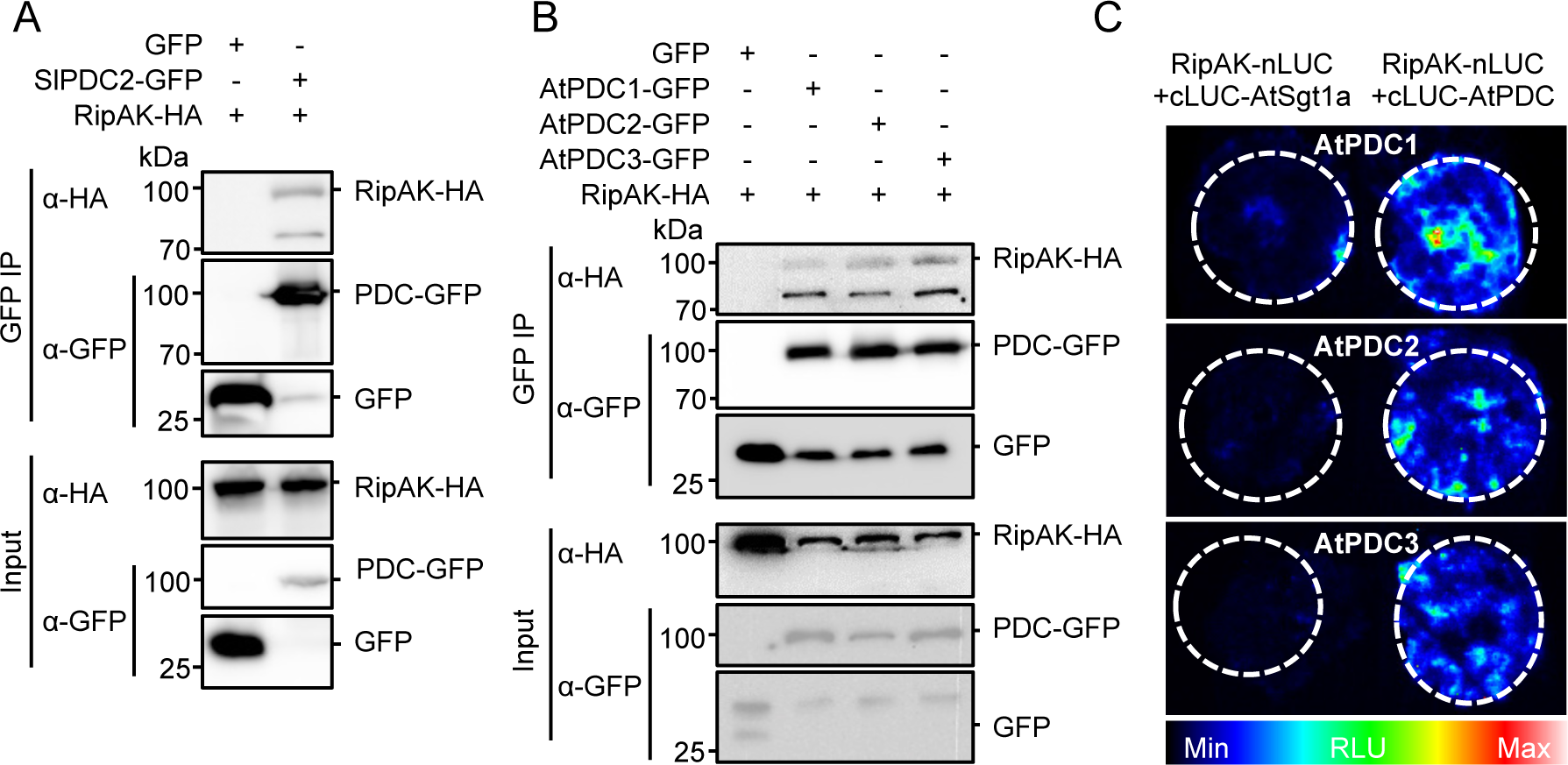
RipAK interacts with pyruvate decarboxylases. (A and B) Co-immunoprecipitation assays to determine interactions between RipAK and PDCs from tomato (A) and Arabidopsis (B). *A. tumefaciens* containing the indicated constructs were inoculated in *N. benthamiana* leaves and samples were taken 44 hours post-inoculation (hpi). Immunoblots were analysed with anti-GFP and anti-HA antibodies, and protein marker sizes are provided for reference. These experiments were performed 3 times with similar results. (C) RipAK interacts directly with Arabidopsis PDCs as determined by Split-LUC assays. RipAK-nLUC and cLUC-AtPDCs were co-expressed in *N. benthamiana* leaves, and luciferase complementation was observed 44 hpi. A colour code representing the relative luminescence is shown for reference. cLUC-AtSgt1a was used as negative interaction control. The accumulation of all the proteins was verified and is shown in Figure S3C.

### *R. solanacearum* infection enhances PDC activity

In conditions of anoxia, plants use PDCs to convert pyruvate into acetaldehyde to contribute to the fermentation process; acetaldehyde can be detoxified into acetate by aldehyde dehydrogenases (Kürsteiner et al., 2003) (Figure S3D). The expression of *PDC* genes is low in basal conditions, but is up-regulated in conditions of anoxia, drought, and other stresses (Kim et al., 2017; Kürsteiner et al., 2003; Mithran et al., 2014). Interestingly, the expression of several *PDC* orthologs in different plant species is up-regulated upon *R. solanacearum* inoculation (Table S2). This prompted us to measure PDC activity in Arabidopsis and tomato during *R. solanacearum* infection. As shown in Figure 3A, Arabidopsis Col-0 WT seedlings showed an increase in PDC activity as early as 2 days after inoculation with *R. solanacearum* GMI1000. A similar pattern was observed in tomato stems starting one day after injection of *R. solanacearum* GMI1000 (Figure 3B). PDC activity did not increase significantly upon inoculation with a non-pathogenic *ΔhrpG* mutant strain (Figure 3A), which cannot express the T3SS and other virulence factors (Valls et al., 2006), is impaired in vascular colonization (Vasse et al., 2000), and does not cause disease symptoms (Brito et al., 1999). It is noteworthy that, during an active infection by a pathogenic strain, *R. solanacearum* rapidly consumes the available oxygen present in the xylem, generating a hypoxic environment (Dalsing et al., 2015). Therefore, our data could suggest that the fast replication of *R. solanacearum* GMI1000 (but not the non-pathogenic mutant) and the subsequent depletion of available oxygen triggers a response to hypoxia in infected tissues, including a rapid increase of PDC activity. However, given the complexity of the *R. solanacearum* infection process, other explanations for this response cannot be ruled out.

**Figure 3.**
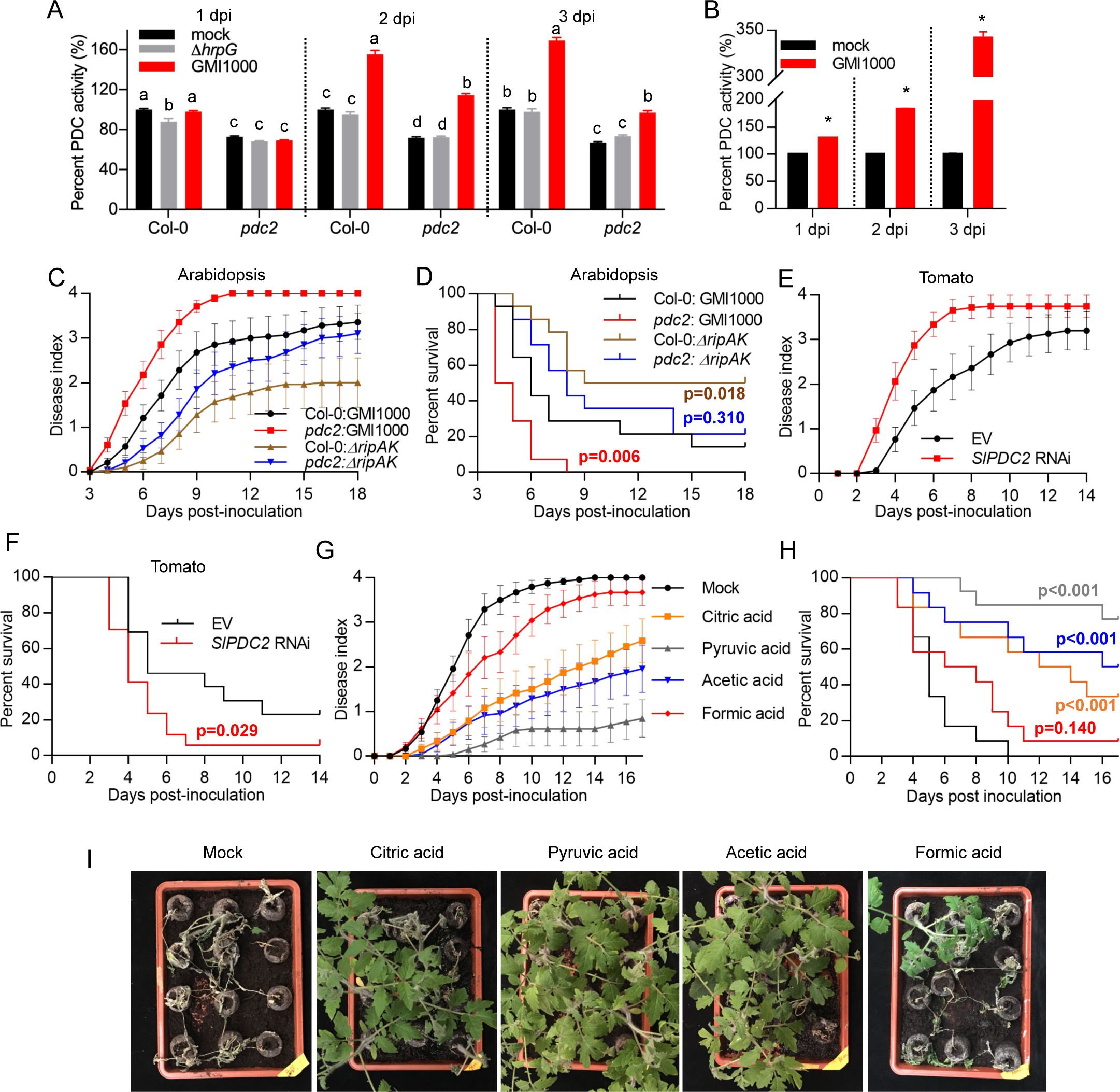
The PDC-mediated pathway contributes to resistance against *R. solanacearum*. (A and B) *R. solanacearum* inoculation in Arabidopsis seedlings (A) or tomato stems (B) stimulates PDC enzymatic activity. (A) Roots of 8 day-old Arabidopsis seedlings were inoculated with 10 µl of a 10^5^ cfu ml^-1^ *R. solanacearum* suspension. GMI1000 WT or a *ΔhrpG* mutant were used, as indicated, and water was used as mock treatment. (B) Stems of 3.5 week-old tomato plants were injected with 5 µl of a 10^5^ cfu ml^-1^ *R. solanacearum* suspension. PDC activity was determined in whole seedlings (A) or stem tissue (B) 1, 2, and 3 dpi, and is represented as percentage PDC activity relative to the wild-type mock control for each day. In A, different letters indicate significantly different values within each time point, as determined using a one-way ANOVA statistical test (p<0.05). In B, asterisks indicate values significantly different to the mock control for each day, as determined using a Student’s t test (p<0.001). Values represent mean ± SEM (n=8). Small error bars may not be visible in some columns. These experiments were performed 3 times with similar results. (C and D) Soil-drenching inoculation assays in Arabidopsis Col-0 WT or *pdc2* mutants, performed with GMI1000 WT or the *ΔripAK* mutant. n≥15 plants per genotype. (E and F) Soil-drenching inoculation assays in tomato plants with transgenic roots expressing an empty vector (EV) or an RNAi construct to silence *SlPDC2*, performed with GMI1000 WT. Transgenic roots were generated using *Agrobacterium rhizogenes* (see methods). n≥8 plants per genotype. (G and H) Soil-drenching inoculation assays in tomato plants upon pre-treatment with a 30 mM solution of the indicated organic acids or water (as mock control). Treatments were performed by placing the pots on a layer of wet towel paper containing the organic acids for 9 days, and then washed and watered normally without treatment for 3 days before inoculation with *R. solanacearum* GMI1000 WT. n≥12 plants per treatment. In C, E, and G the results are represented as disease progression, showing the average wilting symptoms in a scale from 0 to 4 (mean ± SEM). D, F, and H show the survival analysis of the data in C, E, and G, respectively; the disease scoring was transformed into binary data with the following criteria: a disease index lower than 2 was defined as ‘0’, while a disease index equal or higher than 2 was defined as ‘1’ for each specific time point. Statistical analysis was performed using a Log-rank (Mantel-Cox) test, and the corresponding p value is shown in the graph with the same colour as each curve. Four, three, and seven independent biological replicates were performed for inoculations in C, E, and G, respectively, and composite data representations are shown in Figure S5. (I) Representative images of the inoculated plants in G-H 17 dpi.

### PDCs contribute to plant resistance against *R. solanacearum*

PDCs contribute to plant resistance against abiotic stress, including anoxia and drought (Kim et al., 2017; Kürsteiner et al., 2003). In order to determine if PDCs contribute to resistance against *R. solanacearum*, we first ordered mutant lines with T-DNA insertions in *AtPDC1*, *AtPDC2*, and *AtPDC3* (Figure S4A), and determined the expression of these genes in seedlings of each mutant line. Although the expression of each gene was virtually abolished in its respective mutant line, we noticed that the *pdc1* mutant line showed constitutive up-regulation of the expression of the *PDC3* gene, and the *pdc3* mutant line showed constitutive up-regulation of the expression of the *PDC2* gene (Figures S4B-D), which may reflect compensatory effects among functionally redundant genes. On the contrary, the *pdc2* mutant line showed slightly reduced expression of *PDC1* gene. We then analysed PDC activity in each mutant line; although all three mutants showed lower PDC activity compared to WT plants in specific biological replicates, only the *pdc2* mutant displayed a reproducible reduction in all replicates (Figure S4E). Accordingly, the enhancement of PDC activity observed during *R. solanacearum* infection was significantly compromised in *pdc2* mutant plants (Figure 3A). For these reasons, we decided to use the *pdc2* mutant for further experiments. Compared to WT plants, *pdc2* mutants showed enhanced susceptibility to bacterial wilt upon inoculation with *R. solanacearum* GMI1000 (Figures 3C, 3D, S5A, and S5B). Interestingly, the *pdc2* mutation was able to rescue the virulence attenuation caused by the *ΔripAK* mutation (Figures 3C, 3D, S5A, and S5B). These results indicate that PDC2 contributes to resistance against bacterial wilt in Arabidopsis, and point at PDC2 as a relevant target of RipAK virulence activity.

To determine the contribution of PDCs to bacterial wilt resistance in tomato, we used tomato plants with transgenic roots expressing an RNAi construct that silenced the expression of *SlPDC2* (Figure S5C). These plants showed significantly enhanced susceptibility upon inoculation with *R. solanacearum* GMI1000 (Figures 3E, 3F, S5D, and S5E), indicating that PDCs also contribute to resistance against bacterial wilt in tomato. We were unable to generate Arabidopsis plants overexpressing *AtPDC* genes, and tomato plants with roots overexpressing *SlPDC2* showed very strong pleiotropic effects, suggesting that, in our experimental conditions, plants may not be able to tolerate sustained overexpression of *PDC* genes.

We have observed that *R. solanacearum* infection causes a prompt activation of PDC activity, probably as a result of the active bacterial replication and the subsequent oxygen depletion (Figures 3A and 3B). The activation of PDC activity leads to a dynamic metabolic flux conversion from glycolysis into acetate synthesis, conferring tolerance to conditions of low water availability (Kim et al., 2017). The enhanced susceptibility to bacterial wilt symptoms observed in plants with mutated or silenced *PDCs* could suggest that the activation of the PDC-mediated acetate pathway may trigger a response that prepares the plant to face better the water deficiency caused by the vascular clogging associated to *R. solanacearum* infection, which eventually causes disease symptoms. In such scenario, plants with deficient PDC activity may develop faster and stronger symptoms, which is in agreement with our observations in Arabidopsis and tomato (Figures 3 and S5).

### The PDC-mediated pathway contributes to resistance against *R. solanacearum* in tomato

To determine whether the PDC-mediated acetate pathway enhances resistance to *R. solanacearum*, we pre-treated tomato plants with exogenous pyruvic acid and acetic acid, as substrate and product of the pathway, respectively. Treatments were performed by placing the pots on a layer of wet towel paper containing the organic acids for 9 days, as previously described (Kim et al., 2017). The pots were then washed to remove the remaining acids and watered normally without treatment for 3 days before bacterial inoculation. Pre-treatment with both pyruvic and acetic acid strongly enhanced resistance against *R. solanacearum* infection, shown as a drastic reduction and delay of wilting symptoms (Figures 3G-I, S5F and S5G). Pre-treatment with other organic acids caused different outcomes: citric acid significantly reduced disease symptoms (Figures 3G-I), although its impact across multiple independent experiments was not as strong as those of pyruvic or acetic acid (Figures S5F and S5G), and formic acid did not have a significant impact on disease symptoms (Figures 3G-I, S5F and S5G). Interestingly, pre-treatment with pyruvic and acetic acid did not affect *R. solanacearum* replication upon injection in tomato stems (Figure S5H). Considering that a deficiency in the PDC pathway enhances the severity of bacterial wilt (Figure 3A-F), and that pyruvic and acetic acid treatments enhance disease resistance (Figure 3G-I), the activation of the PDC-mediated acetate pathway may indeed contribute to a reduction of disease symptoms by a similar mechanism involved in resistance against drought, although we should not discard the possibility that this pathway actively contributes to resistance against bacterial proliferation, for example, by modulating hormone signalling (Kim et al., 2017).

### RipAK inhibits PDC oligomerisation and activity *in vivo*

Given that the PDC pathway contributes to disease resistance against bacterial wilt and that mutation of *pdc2* (which reduces PDC activity) rescues the virulence attenuation of a *R. solanacearum ΔripAK* mutant (Figure 3), we sought to determine whether RipAK inhibits the enzymatic activity of PDC2. Overexpression of *AtPDC2* in *N. benthamiana* enhanced PDC activity in comparison to control conditions (Figures 4A and 4B). The simultaneous expression of RipAK did not affect the accumulation of AtPDC2 (Figure 4B), but significantly reduced PDC activity (Figures 4A and 4B). PDC enzymes are known to form oligomers, and molecular studies in yeast PDCs have shown that oligomerisation is required for enzymatic activity (Killenberg-Jabs et al., 2001). Upon transient expression in *N. benthamiana*, we also detected direct interaction between different AtPDC2 versions tagged with different halves of luciferase (Figure 4C-E). Interestingly, AtPDC2 oligomerisation *in planta* was inhibited by RipAK (Figure 4C-E), suggesting that this could be the molecular mechanism behind the RipAK-mediated inhibition of PDC activity. It is generally accepted that effector proteins often display multiple targets in plant cells (Macho and Zipfel, 2015), and RipAK has also been shown to target and suppress the activity of catalases in tobacco cells (Sun et al., 2017). Interestingly, like PDCs, catalases are active as oligomers (Nicholls et al., 2000). Although the mechanism of the targeting of catalases is unclear, it is possible that RipAK inhibits the activity of specific host target enzymes during the infection by inhibiting their oligomerisation or their association with interacting partners required for their enzymatic activity.

**Figure 4.**
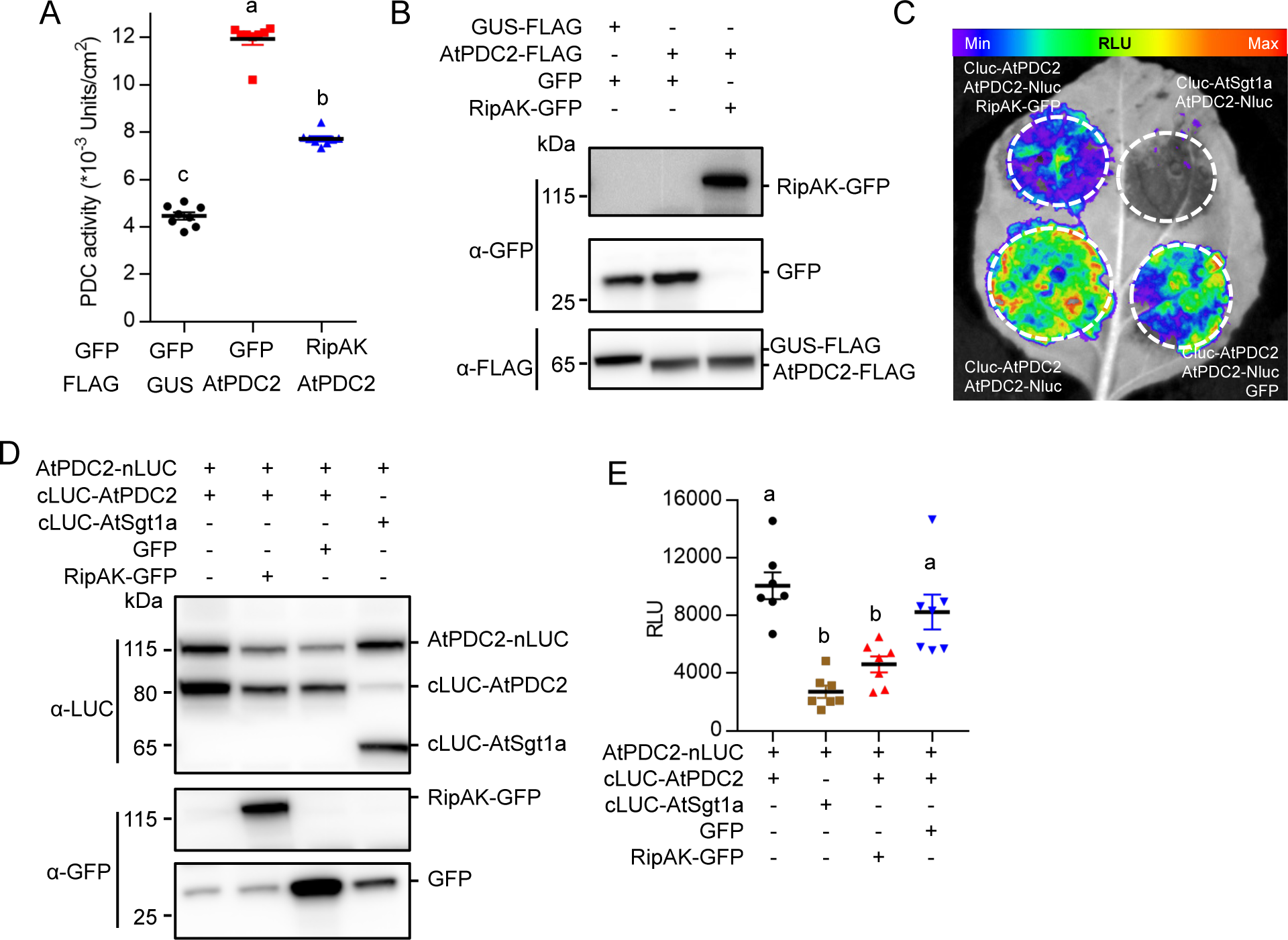
RipAK inhibits PDC oligomerisation and activity *in vivo*. (A) RipAK inhibits AtPDC2 activity in *N. benthamiana.* AtPDC2-FLAG was expressed in *N. benthamiana* leaves using Agrobacterium, and GUS-FLAG was used as control. RipAK-GFP (or GFP, as control) was co-expressed with the FLAG-tagged proteins. PDC activity was determined 36 hpi (mean ± SEM, n=8 per sample), and is represented as units per area of sampled leaf tissue. (B) Protein accumulation in the tissues used to measure PDC activity shown in (A). (C-E) RipAK inhibits AtPDC2 oligomerisation. AtPDC2-nLUC and cLUC-AtPDC2 were co-expressed in *N. benthamiana* leaves to determine AtPDC2 oligomerisation, and AtPDC2-nLUC was co-expressed with cLUC-AtSgt1a as negative control. RipAK-GFP (or GFP, as control) was co-expressed with AtPDC2-nLUC and cLUC-AtPDC2 to determine interference with AtPDC2 oligomerisation. Luciferase complementation was observed 44 hpi, and is shown in (C). A colour code representing the relative luminescence is shown for reference. (D) Protein accumulation in the tissues used for Split-LUC assays. (E) Quantification of luminescence as relative luminescence units (RLU), as detailed in the methods section (mean ± SEM, n=8 per sample). Different letters indicate significantly different values, as determined using a one-way ANOVA statistical test (p<0.05). The immunoblots in this figure were developed using anti-GFP, anti-FLAG, or anti-LUC antibody; the relative position of the different proteins in the blots and protein marker sizes are provided for reference. These experiments were performed 3 times with similar results.

## Conclusions

Bacterial pathogens employ T3Es to suppress immunity and manipulate other cellular functions, including the subversion of plant metabolism by different means (Macho, 2016). Recent studies have revealed that *R. solanacearum* T3Es seems to be particularly prolific at altering plant metabolism upon delivery inside plant cells: RipTPS catalyzes the production of trehalose (Poueymiro et al., 2014), Brg11 induces an increase in polyamine levels, triggering a defence reaction that likely inhibits other microbial competitors (Wu et al., 2019), and RipI induces the production of GABA to support bacterial nutrition (Xian et al., 2019).

Upon invasion of plant tissues, *R. solanacearum* colonizes xylem vessels and replicates rapidly, which depletes the available oxygen (Dalsing et al., 2015). The rapid increase of PDC activity in plant tissues undergoing *R. solanacearum* infection (Figure 3) suggests that plants may respond to pathogen-induced hypoxia by up-regulating *PDC* genes. Subsequently, the activation of the PDC-acetate pathway contributes to alleviating disease-associated wilting symptoms (Figure 3). Given that disease-associated wilting symptoms are likely produced by the restriction in water conductivity derived from vascular clogging, the contribution of the PDC-acetate pathway to disease resistance likely resembles its contribution to drought resistance (Kim et al., 2017), constituting a physiological form of disease resistance. Similarly, ABA, which also acts as a drought stress signal in plants, has been shown to contribute to plant resistance against bacterial wilt (Feng et al., 2012). In addition to this, PDCs may participate in metabolic functions that contribute to the activation of other immune responses. This response, leading to a delay or abolishment of disease symptoms, would also interfere with the bacterial life cycle by impeding bacteria to return to the soil and invade additional plants. *R. solanacearum* may have evolved to counteract such plant response by secreting RipAK, which associates with PDCs and inhibits PDC activity. In agreement with this hypothesis, bacteria lacking RipAK induce slower disease symptoms, while plants with deficient PDC activity develop stronger disease symptoms, partially rescuing the virulence attenuation of a *R. solanacearum ΔripAK* mutant. The virulence activity of RipAK would therefore enable bacteria to complete its life cycle and infect new host plants. Thus, the study of RipAK virulence activity has allowed us to uncover the function of the PDC pathway in disease tolerance, shedding light on the integration between plant responses to biotic and abiotic stresses.

## Acknowledgements

We thank Nemo Peeters and Anne-Claire Cazale for sharing unpublished biological materials, Motoaki Seki and Jian-Min Zhou for sharing biological materials, Rosa Lozano-Duran for critical reading of this manuscript, Xinyu Jian for technical and administrative assistance during this work, and all the members of the Macho and Lozano-Duran laboratories for helpful discussions. We thank the PSC Cell Biology, Proteomics, and Metabolomics core facilities for assistance with enzymatic activity assays, confocal microscopy, and mass spectrometry. This work was supported by the Strategic Priority Research Program of the Chinese Academy of Sciences (grant XDB27040204), the National Natural Science Foundation of China (NSFC; grant 31571973), the Chinese 1000 Talents Program, and the Shanghai Center for Plant Stress Biology (Chinese Academy of Sciences). The authors have no conflict of interest to declare.

## Author contributions

Y.W. and A.P.M. designed the work, supervised experiments, and analysed data. Y.W. performed most of the experimental work. R.J.L.M, G.Y., A.Z., H.X., J.S.R., and Y.S. performed additional experiments. Y.W. and A.P.M. wrote the manuscript with inputs from all the authors.

## MATERIALS AND METHODS

### KEY RESOURCES TABLE

| REAGENT or RESOURCE | SOURCE | IDENTIFIER |
| --- | --- | --- |
| **Antibodies** |  |  |
| Mouse monoclonal anti-GFP | Abiocode | Cat# M0802-3a |
| Mouse monoclonal anti-FLAG | Abmart | Cat# M20008 |
| Rabbit polyclonal anti-Luciferase | Sigma | Cat# L0159 |
| Mouse monoclonal anti-HA | Roche | Cat# 12CA5 |
| anti-Mouse IgG-Peroxidase | Sigma | Cat# A2554 |
| anti-Rabbit IgG-Peroxidase | Sigma | Cat# A0545 |
| **Bacterial and Virus Strains** |  |  |
| *Escherichia coli* DH5a | Transgen | CD501-3 |
| *Agrobacterium tumefaciens* GV3101 | Weidi Bio | AC1001 |
| *Agrobacterium tumefaciens* GV3101 (PMP90RK) | BioRc | Lot#20150202 |
| *Agrobacterium rhizogenes* MSU440 | (Morcillo et al., 2020) | N/A |
| *Ralstonia solanacearum* GMI1000 | (Salanoubat et al., 2002) | N/A |
| *Ralstonia solanacearum* ΔripAK | This work | N/A |
| *Ralstonia solanacearum* ΔripAK/RipAK | This work | N/A |
| **Chemicals, Peptides, and Recombinant Proteins** |  |  |
| Protease Inhibitor Cocktail for plant cell and tissue extracts, DMSO solution | Sigma | P9599 |
| GFP-Trap_A | Chromotek | Cat# gta-100 |
| XenoLight D-Luciferin | PerkinElmer | Cat# 122799 |
| Citric Acid | Sigma | Lot#SLBR376 5V |
| Sodium pyruvate | Sigma | Lot#SLBW601 9 |
| NaDH-Na2 | Yeasen | Lot#N03702 |
| Alcohol Dehydrogenase (ADH) | Sigma | Lot#SLBH097 7V |
| Thiamine Pyrophosphate | Sangon Biotech | Lot#TB0939 |
| Pyruvic Acid | Sangon Biotech | Lot#PD0452 |
| Acetic Acid | Hushi | Lot#M0130-24 81 |
| XenoLight D-Luciferin | PerkinElmer | Cat# 122799 |
| **Critical Commercial Assays** |  |  |
| pENTR/D-TOPO Cloning Kit | Invitrogen | Cat#<br>K240020SP |
| Gateway LR Clonase II Enzyme Mix | Invitrogen | Cat#<br>11791100 |
| Experimental Models: Organisms/Strains |  |  |
| Arabidopsis: *pd<sup>c</sup>1* | (Gravot et al., 2016; Stepanova et al., 2011) | N/A |
| Arabidopsis: *pd<sup>c</sup>2* | (Gravot et al., 2016; Stepanova et al., 2011) | N/A |
| Arabidopsis: *pd<sup>c</sup>3* | (Stepanova et al., 2011) | N/A |
| *Solanum lycopersicum* cv. Moneymaker | N/A | N/A |
| Primers see Table S3 |  |  |
| Recombinant DNA |  |  |
| pENTR/D-TOPO | Invitrogen | Cat#<br>K240020SP |
| pXCSG-HA Strep | (Witte et al., 2004) | N/A |
| pGWB505 | (Nakagawa et al., 2007b) | N/A |
| pGWB511 | (Nakagawa et al., 2007b) | N/A |
| pGWB554 | (Nakagawa et al., 2007b) | N/A |
| pGWB-nLUC | (Wang et al., 2019b) | N/A |
| pGWB-cLUC | (Yu et al., 2019b) | N/A |
| pEASYBLUNT-LB-Gm-RB | This work | N/A |
| pRCT-pRipAK-RipAK | This work | N/A |
| pGWB505-RipAK-GFP | This work | N/A |
| pGWB-RipAK-nLUC | This work | N/A |
| pGWB-cLUC-AtPDC1 | This work | N/A |
| pGWB-cLUC-AtPDC2 | This work | N/A |
| pGWB-cLUC-AtPDC3 | This work | N/A |
| pGWB-AtPDC2-nLUC | This work | N/A |
| pXCSG-HA Strep-RipAK | This work | N/A |
| pXCSG-HA Strep-GFP | (Sang et al., 2016) | N/A |
| pGWB505-AtPDC1-GFP | This work | N/A |
| pGWB505-AtPDC2-GFP | This work | N/A |
| pGWB505-AtPDC3-GFP | This work | N/A |
| pGWB505-SIPDC2-GFP | This work | N/A |
| pGWB511-AtPDC2-FLAG | This work | N/A |
| pGWB511-GUS-FLAG | (Yu et al., 2019a) | N/A |
| pGWB505-GFP | (Sang et al., 2016) | N/A |
| pGWB554-AtPDC1-RFP | This work | N/A |
| pGWB554-AtPDC2-RFP | This work | N/A |
| pGWB554-AtPDC3-RFP | This work | N/A |
| pGWB-cLUC-AtSGT1a | (Yu et al., 2019b) | N/A |
| pK7GWIWG2_II-RedRoot | (Morcillo et al., 2020) | N/A |
| pK7GWIWG2_II-RedRoot-SIPDC2 | This work | N/A |
| pMD1 | (Li et al., 2013) | N/A |
| pMD1-PTS1- mTurquoise2 | This work | N/A |
| Software and Algorithms |  |  |
| Prism 7 | GraphPad Software | https://www.graphpad.com/scientific-software/prism/ |
| Scaffold 4.0 | Proteome Software | http://www.proteomesoftware.com/products/scaffold/ |
| ImageJ | NIH ImageJ | https://imagej.nih.gov/ij/ |
| Mega X | Mega X | https://megasoftware.net/ |

### CONTACT FOR REAGENT AND RESOURCE SHARING

Further information and requests for resources and reagents should be directed to the Lead Contact, Alberto P. Macho.

### EXPERIMENTAL MODELS AND SUBJECT DETAILS

#### Arabidopsis thaliana

*Arabidopsis thaliana* (Arabidopsis) plants used in this work were in Columbia (Col-0) background. The *pdc1* (SALK090204C) (Gravot et al., 2016; Stepanova et al., 2011), *pdc2* (CS862662) (Gravot et al., 2016; Stepanova et al., 2011), and *pdc3* (SALK087974) (Stepanova et al., 2011) mutant lines were obtained from the Nottingham Arabidopsis Stock Centre. Primers used to genotype these mutants are shown in Table S3 and their target locations in the genes are shown in Figure S4. All the experiments were performed with homozygous plants. Plants used for harvesting seeds were grown on soil in a growth chamber at 23°C, 16 h light/8 h dark, and 70% relative humidity. For PDC enzymatic analysis, seeds were germinated on solid 1/2 Murashige and Skoog (MS) medium and seedlings were grown for 10 days in a long day growth room with 23°C, 16 h light/8 h dark and 70% relative humidity. For *Ralstonia solanacearum* soil drenching assays, seeds were germinated on solid 1/2 MS medium, and seedlings were grown for 1 week before being transferred to Jiffy pots (Jiffy International, Norway). Plants were then grown for 3-4 weeks in a short day growth chamber at 23°C, 12 h light/12 h dark, and 70% relative humidity. After *R. solanacearum* inoculation, plants were transferred to a long day growth chamber at 28°C, 16 h light/8 h dark, and 75% relative humidity.

#### Nicotiana benthamiana

*Nicotiana benthamiana* plants were cultivated in a growth room at 23°C, 16 h light/8 h dark, and 70% relative humidity. Four-week-old *N. benthamiana* plants were used for transient expression and subsequent assays.

#### Solanum lycopersicum

Tomato (*Solanum lycopersicum* cv. Moneymaker) plants were grown in a long day growth chamber at 28°C, 16 h light/8 h dark, and 65% relative humidity. Seeds were germinated on soil for 10 days and then transferred to Jiffy pots for further treatment with organic acids or *R. solanacearum* inoculation. After *R. solanacearum* inoculation, plants were transferred to a long day growth chamber at 28°C, 16 h light/8 h dark, and 75% relative humidity.

#### Bacterial strains

*Agrobacterium tumefaciens* GV3101 or GV3101 (PMP90RK) was used for transient expression in *N. benthamiana*. *Agrobacterium rhizogenes* MSU440 for expression in tomato roots. *A. tumefaciens* strains were grown on solid LB medium plates with the appropriate antibiotics for 2 days at 28 °C, and then inoculated in liquid LB medium with appropriate antibiotics to grow overnight at 28 °C. The antibiotic concentrations used were 25 µg mL^-1^ rifampicin, 50 µg mL^-1^ gentamicin, 50 µg mL^-1^ kanamycin, 50 µg mL^-1^ spectinomycin, and 50 µg mL^-1^ carbenicillin. *R. solanacearum* strains were grown in the same conditions using BG medium. (Plener et al., 2012)

## METHODS DETAILS

### Generation of plasmid constructs, transgenic plants, and *R. solanacearum* mutant strains

The RipAK coding region (Rsc2359) in pDONR207 (a gift from Anne-Claire Cazale and Nemo Peeters, LIPM, Toulouse, France) was used as a template to amplify the sequence encoding the full RipAK or the truncated version lacking the 70 N-terminal amino acids (primers are detailed in Table S3). Fragments were cloned into pENTR/D-TOPO (Thermo Scientific, USA) and then subcloned into the expression vectors pGWB505 (Nakagawa et al., 2007a), pXCSG-HAStrep (Witte et al., 2004), and pGWB-cLUC/nLUC (Wang et al., 2019a; Yu et al., 2019b) via attL-attR recombinant (LR) reactions (Thermo Scientific, MA, USA). The full length of mTurquoise2 fused to a PTS1 was amplified from the pmTurquoise2-Peroxi vector (Goedhart et al., 2012) using the primers listed in Table S3. The amplified fragment was cloned into the *Not*I/*Asc*I sites of pENTR-D and then subcloned into the expression vector pMD1 (Li et al., 2013). Arabidopsis *AtPDC1* (AT4G33070), *AtPDC2* (At5G54960), and *AtPDC3* (At5G01330) were amplified from cDNA of Arabidopsis Col-0 using the primers detailed in Table S3, cloned into pENTR-D/TOPO, and then subcloned into the expression vectors pGWB505, pGWB554, pGWB511 (Nakagawa et al., 2007a), and pGWB-cLUC/nLUC. To generate the *R. solanacearum ΔripAK* mutant strain, the RipAK gene was replaced by a gentamicin resistance gene as described by Zumaquero (Zumaquero et al., 2010). The RipAK flanking regions, left border (LB) and right border (RB), were amplified by PCR and recombined into pEASYBLUNT vector; subsequently, a gentamicin resistance cassette was inserted between LB and RB through *Eco*R I digestion and T4 ligation, resulting in pEASYBLUNT-LB-Gm-RB. The pEASYBLUNT-LB-Gm-RB plasmid was introduced into *R. solanacearum* GMI1000 strain by natural transformation (González et al., 2011). The *ΔripAK* mutant strain was selected with 10 µg mL^-1^ gentamicin and confirmed using RipAK specific primers (Table S3). To generate the *ΔripAK/RipAK* complementation strain, the RipAK gene (including 253bp upstream of RipAK gene start codon ATG) was cloned into pENTR-D/TOPO, introduced into pRCT-GWY vector by LR reaction, and then transformed into the *ΔripAK* mutant strain (Henry et al., 2017). The *ΔripAK/RipAK* complementation strain was selected with 10 µg mL^-1^ tetracycline and confirmed using RipAK specific primers (Table S3).

### Pathogen inoculation assays

For *R. solanacearum* soil drenching inoculation, 4.5-week old Arabidopsis (at least 20 plants per genotype) or 3.5-week old tomato plants (at least 12 plants per genotype) were used (the exact number for each experiment is indicated in the figure legend). Plants grown in Jiffy pots were inoculated by soil drenching with a bacterial suspension containing 10^8^ colony-forming units per mL (CFU mL^-1^). 30 mL of inoculum of each strain was used to soak each plant. After a 20-minute incubation with the bacterial inoculum, plants were transferred from the bacterial solution to a bed of potting mixture soil in a new tray (Vailleau et al., 2007). Scoring of visual disease symptoms on the basis of a scale ranging from ‘0’ (no symptoms) to ‘4’ (complete wilting) was performed as previously described (Vailleau et al., 2007). To perform survival analysis, the disease scoring was transformed into binary data with the following criteria: a disease index lower than 2 was defined as ‘0’, while a disease index equal or higher than 2 was defined as ‘1’ for each specific time point (days post-inoculation, dpi) (Remigi et al., 2011).

Stem injection assays were performed as previously described (Yu et al., 2019b). Briefly, 5 µL of a 10^6^ CFU mL^-1^ bacterial suspension was injected into the stems of 4-week-old tomato plants and 2.5 µL of xylem sap was collected from each plant for bacterial number quantification at the indicated times. Injections were performed 2 cm below the cotyledon emerging site in the stem, and the samples were taken at the cotyledon emerging site.

To measure PDC activity upon bacterial inoculation, Arabidopsis seedlings were grown on 1/2 MS solid medium plates for one week. Seedlings were inoculated by placing 10 µL of a bacterial inoculum containing 10^5^ CFU mL^-1^ of *R. solanacearum* inoculation on the root tip of each seedling. Seedlings were collected 1, 2, or 3 dpi for PDC activity measurement. Tomato plants were inoculated by stem injection as described above, and samples for PDC enzymatic assay were taken and frozen in liquid nitrogen at 1, 2, and 3 dpi.

### RNAi in tomato roots

To generate the *SlPDC2* RNAi construct, a 204bp fragment of tomato *SlPDC2* (*Solyc02g077240*) was amplified from cDNA of tomato cv. Moneymaker using the primers detailed in Table S3, cloned into pENTR-D/TOPO, and then subcloned into the expression vector pK7GWIWG2_II-RedRoot (http://gateway.psb.ugent.be). The *SlPDC2* cloned fragment shares 81%, 81%, and 84% homology to the respective fragments of *Solyc09g005110*, *Solyc06g082130,* and *Solyc10g076510*, respectively, which are also annotated as SlPDCs (Figure S3).

The generation of tomato plants with transgenic roots was performed as previously described (Morcillo et al., 2020). Briefly, the radicles of tomato seedlings were cut, and the resulting hypocotyls were dipped in *Agrobacterium rhizogenes* MSU440 containing pK7GWIWG2_II-RedRoot::*SlPDC2* or pK7GWIWG2_II-RedRoot (used as control). The seedlings where then incubated to allow the growth of transgenic roots. Three weeks after transformation, seedlings were transferred to Jiffy pots, and soil-drenching inoculation with *R. solanacearum* (OD_600_ of 0.1) was performed three-to-four weeks later as described above. Symptoms were scored as described above. The efficiency of the *SlPDC2* silencing was determined by qRT PCR, and shown in the figure S5.

### Transient expression in *N. benthamiana*

Transient expression in *N. benthamiana* was performed as previously described (Sang et al., 2016). Briefly, *A. tumefaciens* strains carrying the indicated constructs were infiltrated into leaves of 4.5-week-old *N. benthamiana* using an OD_600_ of 0.5. To prepare the inoculum, *A. tumefaciens* was incubated in infiltration buffer (10 mM MgCl_2_, 10 mM MES pH 5.6, and 150 µM acetosyringone) for 2 h before infiltration. The constructs used as controls for transient expression in *N. benthamiana* were: GFP (Sang et al., 2016), cLUC-AtSGT1a and GUS-FLAG (Yu et al., 2019a).

### Confocal microscopy

Confocal microscopy was performed as previously described (Wang et al., 2019a). Briefly, to determine the subcellular localization of tagged proteins, leaf discs were collected from *N. benthamiana* leaves 2 dpi with *A. tumefaciens*, and observed using a Leica TCS SP8 (Leica, Germany) confocal microscope with the following excitation wavelengths: GFP, 488 nm; RFP, 561 nm; Turquoise, 442 nm, and the respective emission wavelengths: GFP, 500-550nm; RFP, 580-610; Turquoise, 455-490 nm.

### Protein extraction and western blots

Protein extraction and western blots were performed as previously described (Sang et al., 2016) with several modifications. Briefly, plant tissues were collected into 2 mL tubes with metal beads and frozen in liquid nitrogen before grinding using a tissue lyser (Qiagen, Germany) for 1 min at 25 rpm/s. Proteins were then extracted using protein extraction buffer (100 mM Tris-HCl, pH7.5; 10% glycerol; 1% NP40, 5 mM EDTA; 5 mM DTT, 1% Protease inhibitor cocktail, 2 mM PMSF, 10 mM sodium molybdate, 10 mM sodium fluoride, 2 mM sodium orthovanadate) and incubated for 10 min on ice. After centrifugation (10 min; 16,000 g), the supernatants were mixed with SDS loading buffer, denatured at 70 °C for 20 min, and resolved using SDS-PAGE. Proteins were transferred to a PVDF membrane and monitored by western blot using the antibodies indicated in KEY RESOURCES TABLE.

### Immunoprecipitation

Co-immunoprecipitation assays were performed as previously described (Sang et al., 2016) with several modifications. Briefly, *N. benthamiana* leaves were infiltrated with *A. tumefaciens* containing the indicated constructs. Total proteins (0.75 g tissue per sample) were extracted as indicated above and immunoprecipitation was performed with 15 µL of GFP-trap beads (ChromoTek, Germany) during a 1-hour incubation at 4 °C. Beads were washed 4 times with wash buffer containing 0.2% NP40. The proteins were stripped from the beads by heating in 30 µL Laemmli buffer for 20 minutes at 75 °C. The immunoprecipitated proteins were separated on SDS-PAGE gels for western blot analysis with the indicated antibodies. The LC-MSMS analysis of immunoprecipitated RipAK-GFP was performed as previously described (Sang et al., 2016)(Sang et al, 2016).

### Split-LUC analysis

Split-LUC assays were performed as previously described (Chen et al., 2008; Wang et al., 2019a) with several modifications. Briefly, *A. tumefaciens* strains containing the indicated constructs were infiltrated into *N. benthamiana* leaves. A construct containing cLUC-AtSgt1a (Yu et al., 2019b) was used as negative control. Split-LUC assays were performed 44 hours post-inoculation (hpi) for RipAK-PDC interaction or 40 hpi for PDC oligomerisation. For CCD imaging, the leaves were infiltrated with 0.1 mM luciferin in water and kept in the dark for 5 min to reduce the background signal before the analysis. The images were taken with either Lumazone 1300B (Scientific Instrument, USA) or NightShade LB 985 (Berthold, Germany). Image J software was used to quantify the luciferase signal. The protein accumulation was determined by immunoblot as described above.

### RNA extraction and quantitative RT-PCR

For RNA extraction, plant tissues were collected in 1.5 mL microfuge tubes with one metal bead and the tubes were immediately placed into liquid nitrogen. Samples were ground thoroughly using a tissue lyser for 1 minute, and placed back in liquid nitrogen. Total RNA was extracted with the E.Z.N.A. Plant RNA kit (Biotek, China) without DNA digestion according to the manufacturer’s manual. RNA samples were quantified with Nanodrop spectrophotometer (ThermoFisher, USA). The first strand cDNA was synthesized with the iScript gDNA Clear cDNA Synthesis Kit (Bio-Rad) using 1 µg RNA. Quantitative RT-PCR (RT-qPCR) was performed using the iTaqTM Universal SYBR Green Supermix (Bio-Rad, USA) and CFX96 Real-time system (Bio-Rad, USA). Primers are listed in Table S3.

### Yeast two-hybrid

Yeast two-hybrid screening was performed by Hybrigenics Services (Evry, France). The coding sequence of full-length RipAK was PCR-amplified and cloned into pB29 as an N-terminal fusion to LexA (RipAK-LexA). The construct was checked by sequencing the entire insert and used as a bait to screen a random-primed tomato roots (*R. solanacearum* and *Meloidogyne incognita*) cDNA library constructed into pP6. pB29 and pP6 derive from the original pBTM116 (Béranger et al., 1997; Vojtek and Hollenberg, 1995) and pGADGH (Bartel et al., 1993) plasmids, respectively. 90 million clones (9-fold the complexity of the library) were screened using a mating approach with YHGX13 (Y187 ade2-101::loxP-kanMX-loxP, matα) and L40ΔGal4 (mata) yeast strains as previously described (Fromont-Racine et al., 1997). 167 His+ colonies were selected on a medium lacking tryptophan, leucine, and histidine. The prey fragments of the positive clones were amplified by PCR and sequenced at their 5’ and 3’ junctions. Only high-confidence clones were considered. The resulting sequences were used to identify the corresponding interacting proteins in the GenBank database (NCBI) using a fully automated procedure.

### PDC activity measurements

PDC activity was determined as described by Boeckx (Boeckx et al., 2017). Plant tissues were ground and homogenized in extraction buffer containing 100 mM 2-(N-morpholino) ethane sulfonic acid (MES) buffer (pH 7.5), 5 mM dithiothreitol (DTT) and 2.5% (w/v) polyvinylpyrrolidone (PVP) and 0.02% (w/v) Triton X-100. For Arabidopsis seedlings, fresh tissues were weighed before being frozen in liquid nitrogen. Samples were ground using a tissue lyser and the extraction was performed using a proportion of 3:1 (v/w) to plant tissue. For *N. benthamiana* leaf tissue, 30 leaf discs per sample (7 mm diameter each) were collected, and 700 µL of extraction buffer were added to each sample. Plant crude extracts were incubated at 4°C for 15 minutes, and then were centrifuged at 16000 g for 20 minutes. Then, 50 µL of the supernatants were added to 150 µL of enzymatic analysis buffer, containing 10 mM MES buffer (pH 6.5), 10 µL 50 µM thiamine pyrophosphate (TPP), 50 mM magnesium chloride (MgCl_2_), 50 Units commercial Alcohol Dehydrogenase (ADH) solution (Sigma, USA), 50 mM sodium pyruvate, and 0.8 mM NADH (Yeasen, China). Samples and buffer were mixed in 96-well transparent plates, 8 technical replicates were performed for each sample, and the oxidation of NADH was measured by continuously recording the decrease in absorbance at 340 nm using a Varioskan flash microplate luminescence reader (ThermoFisher, Germany) at 37°C for 90 minutes (1 measurement per minute). Within 90 minutes, the decrease of NADH usually reached a steady basal level. Enzymatic activity was calculated using the data corresponding to the linear section of the curve as described by Boeckx (Boeckx et al., 2017).

### Treatments with organic acids

Organic acid treatments were performed as previously described (Kim et al., 2017), with several modifications. Briefly, two week-old tomato plants grown on Jiffy pots were pre-treated with 30 mM citric acid, pyruvic acid, acetic acid, or formic acid using 10 mL per plant every day for 9 days. The organic acids were applied by soaking a paper towel located below the Jiffy pots, so that they were absorbed by capillarity. After 9 days, Jiffy pots were washed with water gently without damaging the roots for several times to remove the remaining acids from the soil, and plants were watered without organic acids for 3 days before inoculation with *R. solanacearum*.

### Sequence analysis

The PDC protein sequences from different plant species were retrieved from NCBI (https://www.ncbi.nlm.nih.gov), SolGenomics (https://solgenomics.net) and TAIR (www.arabidopsis.org). To generate the phylogenetic tree, PDC protein sequences were aligned using MEGA X software, using the Maximum likelihood computation method.

### Statistical analysis

Statistical analyses were performed with the Prism 7.0 software (GraphPad). The data are presented as mean ± SEM. The statistical analyses used are described in the figure legends.

---

**Figure S1.**
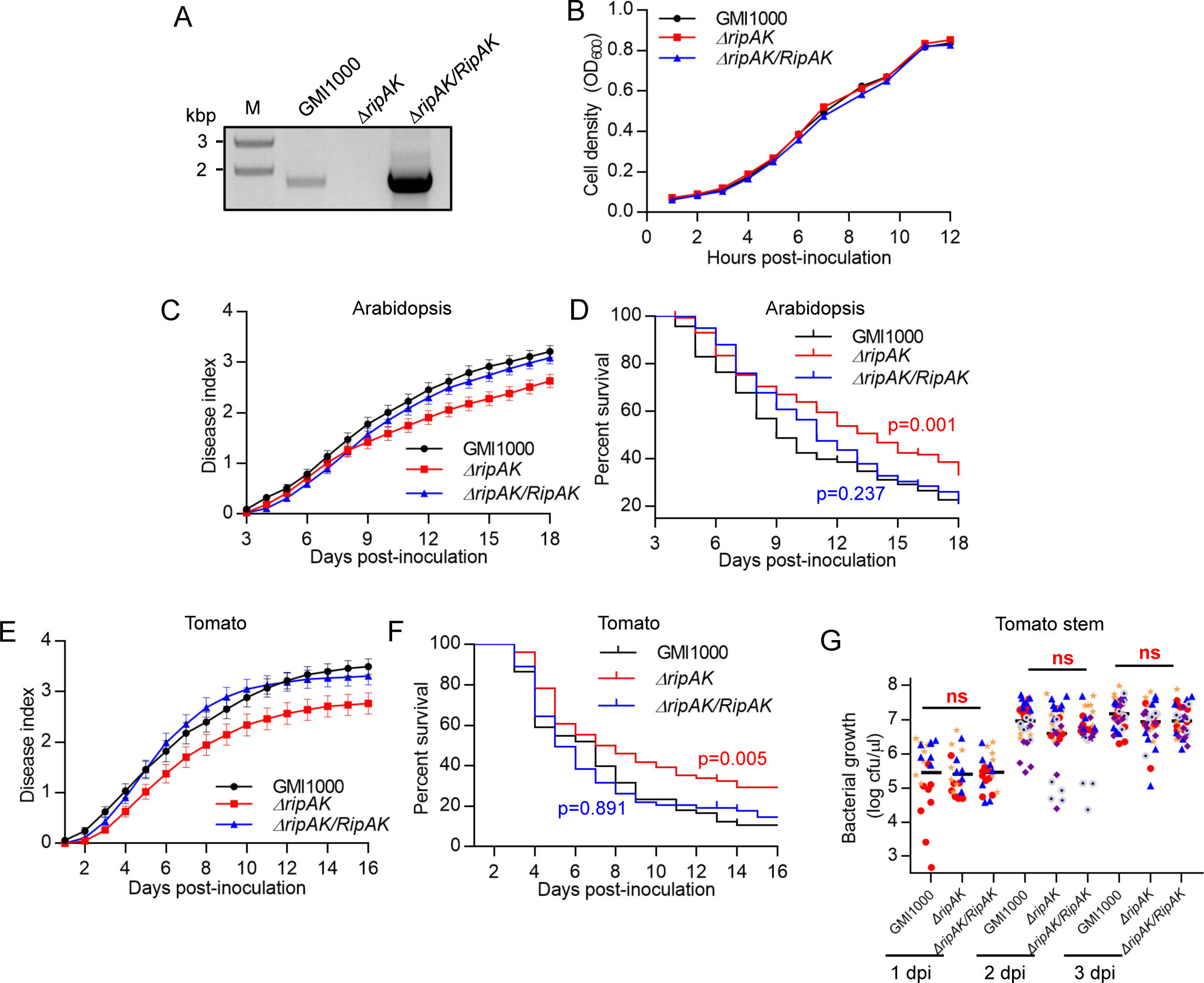
Validation of Δr*ipAK* mutant strains and associated virulence analysis. (A) Genotyping of the Δr*ipAK* mutant and Δr*ipAK/RipAK* complementation strains, using GMI1000 as control. The PCR shows the presence/absence of the *ripAK* fragment in these strains. (B) The Δr*ipAK* mutant and Δr*ipAK/RipAK* complementation strains do not show differences in fitness compared to GMI1000 in nutrient-rich liquid medium. The different strains were inoculated in liquid Phi medium with an initial concentration of OD_600_=0.02, and optical density was measured over time. Values represent mean ± SEM (n=3). (C and D) RipAK contributes to *R. solanacearum* infection in Arabidopsis. Composite data from 9 independent biological repeats (a representative assay is shown in Figure 1A and 1B). All values were pooled together and represented as disease index (C) or percent survival (D). Disease index values represent mean ± SEM (n=158). (E and F) RipAK contributes to *R. solanacearum* infection in tomato. Composite data from 5 independent biological repeats (a representative assay is shown in Figure 1C and 1D). All values were pooled together and represented as disease index (E) or percent survival (F). Disease index values represent mean ± SEM (n=78). Statistical analysis was performed using a Log-rank (Mantel-Cox) test, and the corresponding p value is shown in the graph with the same colour as each curve. (G) The Δr*ipAK* mutant and Δr*ipAK/RipAK* complementation strains do not show differences in growth upon tomato stem injection compared to GMI1000. 3.5-week old tomato plants were injected with 5 µL of a 10^6^ cfu mL^-1^ and samples were collected 1, 2, and 3 dpi. Five independent biological repeats were performed (n=6 plants per strain in each replicate) with similar results. Values from all the replicates are represented in this graph; values with the same colour correspond to the same repeat. ns indicates no significant differences among these strains according to a Student’s t test (p>0.05).

**Figure S2.**
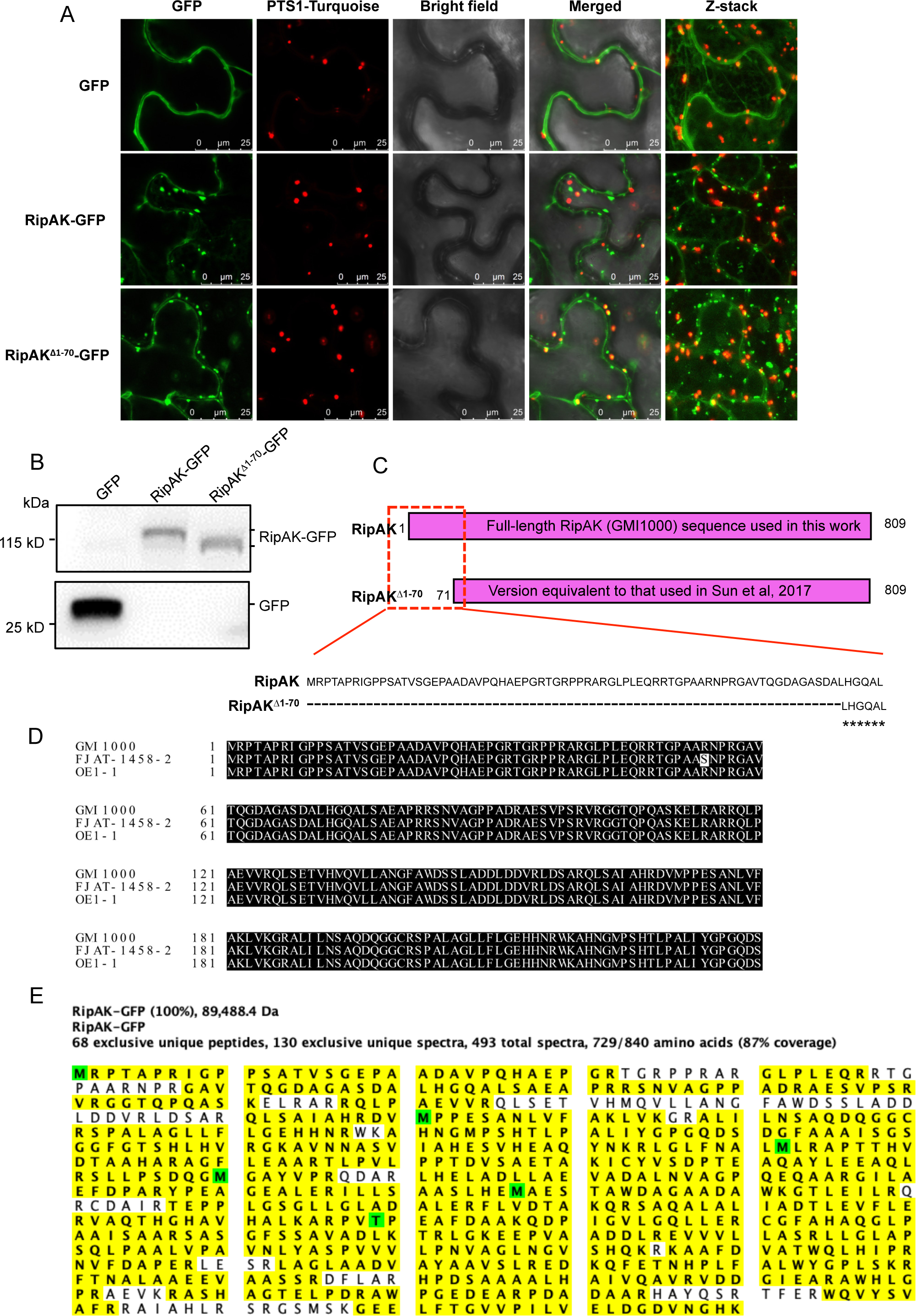
Comparison between the full RipAK reference sequence and the RipAK^Δ1-70^ truncated version. (A) Subcellular localization of RipAK-GFP, RipAK^Δ1-70aa^, and free GFP (as control) in *N. benthamiana* leaf cells observed using confocal microscopy upon transient expression using *A. tumefaciens*. GFP-tagged proteins were co-expressed with PTS1 (peroxisome targeting signal 1) fused to Turquoise fluorescent protein to allow for visualization of peroxisomes. Bright field is provided for reference, and merged signals show the relative localization of GFP and peroxisomes-tagged proteins. Fluorescence was visualized 48 hours-post inoculation. Scale bar = 25 µm. Z-stack shows a vertical cross-section through the observed cells. (B) Western blot to determine the accumulation of GFP tagged proteins in the tissues used for confocal microscopy in (A). Samples were taken 40 hpi, immunoblots were analysed with an anti-GFP antibody, and protein marker sizes are provided for reference. (C) Diagram comparing the full RipAK version used in this work and the truncated version used in Sun et al, (2017). (D) Amino acid sequence of RipAK from different sequenced strains belonging to the phylotype I, including the reference strain GMI1000 (sequence used in this work), showing that the first 70 amino acids are present and highly conserved in different phylotype I strains. Reference sequences were retrieved from the RalstoT3E database (Peeters et al, 2013; Sabbagh et al, 2019; https://iant.toulouse.inra.fr/bacteria/annotation/site/prj/T3Ev3/). (E) The full RipAK-GFP accumulates in *N. benthamiana* tissues upon transient expression using Agrobacterium. Liquid chromatography and Mass spectrometry (LC-MS) analysis was performed after GFP immunoprecipitation. The highlighted tryptic peptides were detected, representing 87% coverage of the total RipAK sequence, including peptides within the first 70 amino acids. Non-highlighted residues represent peptides that were not detected, probably due to technical reasons associated to the tryptic digestion or the LC-MS analysis.

**Figure S3.**
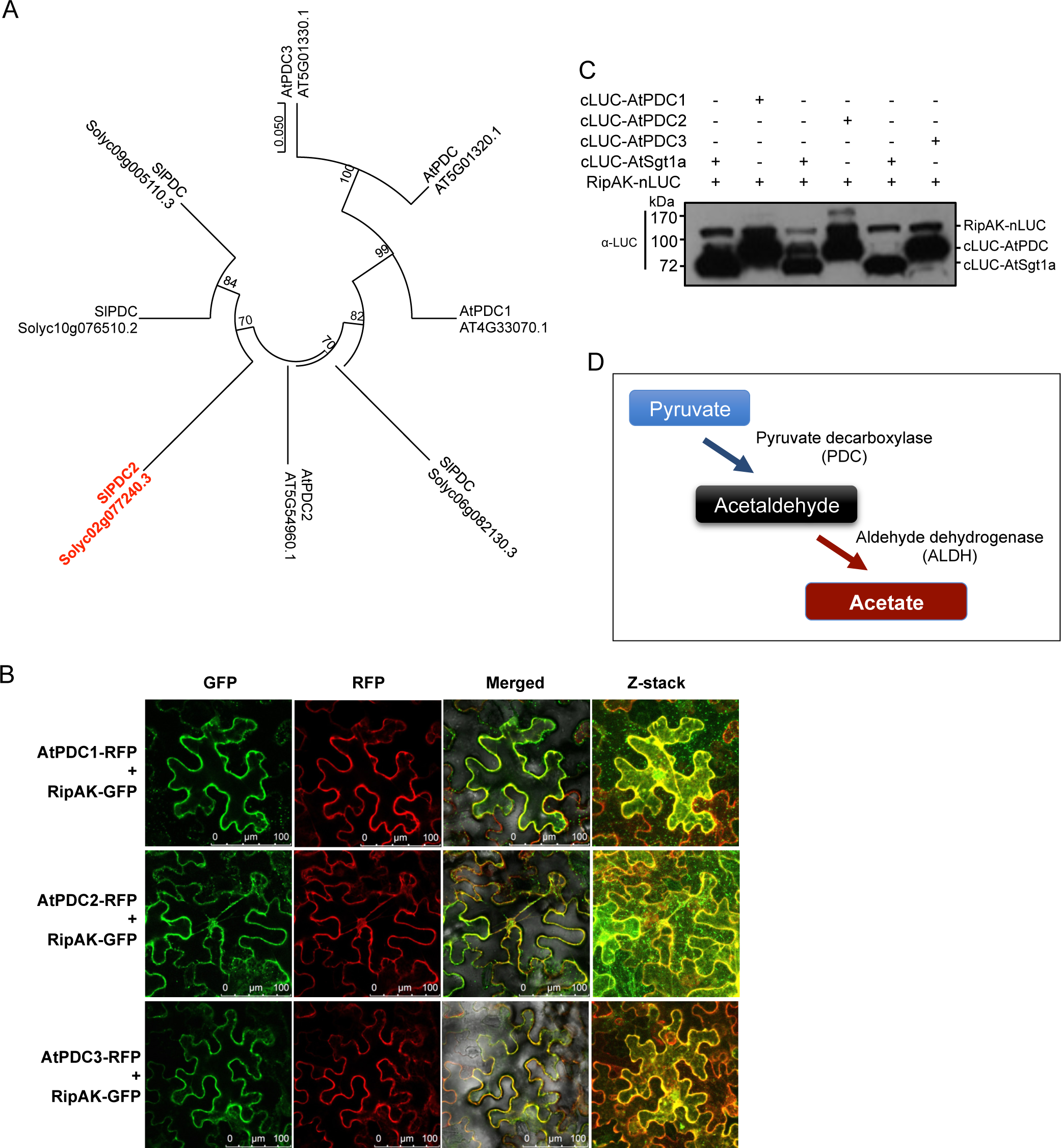
RipAK interacts with PDCs. (A) Phylogenetic tree of PDC proteins from Arabidopsis and tomato. Proteins annotated as “pyruvate decarboxylase” or PDC-family proteins (such as AT5G01320.1) are shown. The SlPDC identified as RipAK interactor (Solyc02g077240) was annotated in this work as SlPDC2 given its high similarity with AtPDC2. The phylogenetic tree was generated using the MEGA X software using the Maximum likelihood method. The percentage of trees in which the associated taxa clustered together is shown next to the branches. (B) Co-localization of RipAK-GFP and AtPDCs tagged with a red fluorescent protein (RFP) in *N. benthamiana* leaf cells observed using confocal microscopy upon transient expression using *A. tumefaciens*. Merged signals show the relative localization of GFP and RFP-tagged proteins. Fluorescence was visualized 40 hpi. Scale bars = 100 µm. Z-stack shows a vertical cross-section through the observed cells. (C) Protein accumulation in the tissues used to perform the Split-LUC assays shown in Figure 2C. RipAK-nLUC and cLUC-AtPDCs were co-expressed in *N. benthamiana* leaves, and cLUC-AtSgt1a was used as negative interaction control. The immunoblot was developed using anti-LUC antibody; the relative position of the different proteins in the blot and protein marker sizes are provided for reference. (D) Simplified diagram of the PDC pathway in stress conditions.

**Figure S4.**
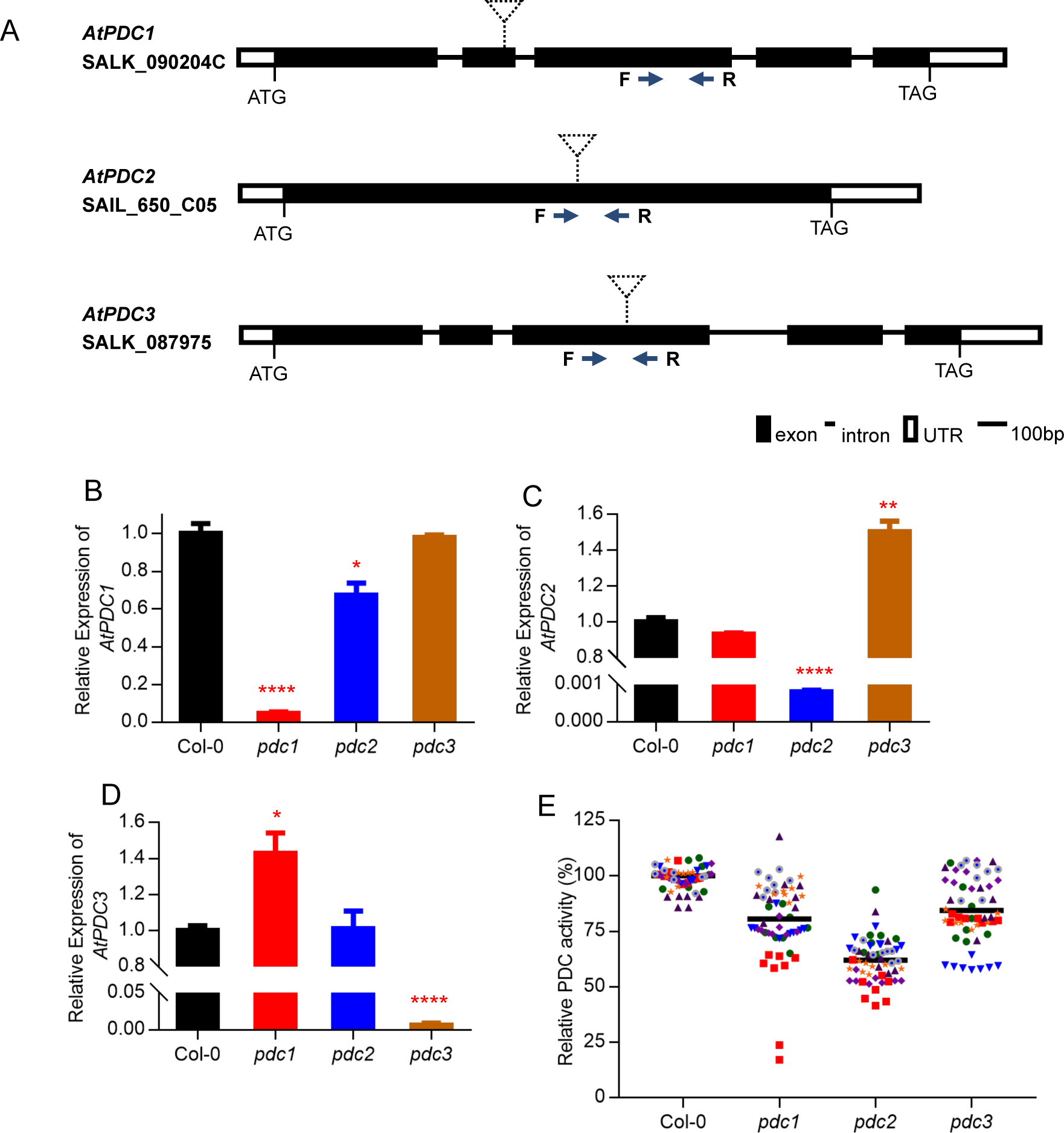
Characterization of Arabidopsis *pdc* mutant lines. (A) Diagram showing the gene structure of *AtPDC1*, *AtPDC2* and *AtPDC3*. Start (ATG) and stop codons are indicated; black boxes represent coding regions, white boxes represent untranslated regions, lines represent intros, and dotted triangles show the location of the T-DNA insertions in each mutant line. F and R indicate the matching sequence of the forward and reverse primers, respectively, used for the subsequent qPCRs to determine gene expression. (B-D) Expression of *AtPDC1, AtPDC2,* and *AtPDC3* in *pdc* mutant lines. Values were normalized to the expression of the AtACT2 gene (AT3G18780) and are shown relative to the expression of each *PDC* gene in Col-0 WT. Values represent mean ± SEM (n=3). The experiments were performed 3 times with similar results. (E) Measurement of PDC activity in Arabidopsis *pdc* mutant lines, using 10 day-old seedlings. Seven independent biological repeats were performed (n=8 in each biological repeat). Values from all the repeats are represented in this graph as percentage of the PDC activity observed in Col-0 WT seedlings in each repeat; values with the same colour correspond to the same repeat. Black bars represent the average values for each mutant. Although *pdc1* and *pdc3* mutants showed reduction in PDC activity in several repeats, only *pdc2* mutant seedlings showed lower PDC activity than Col-0 WT seedlings in all the repeats.

**Figure S5.**
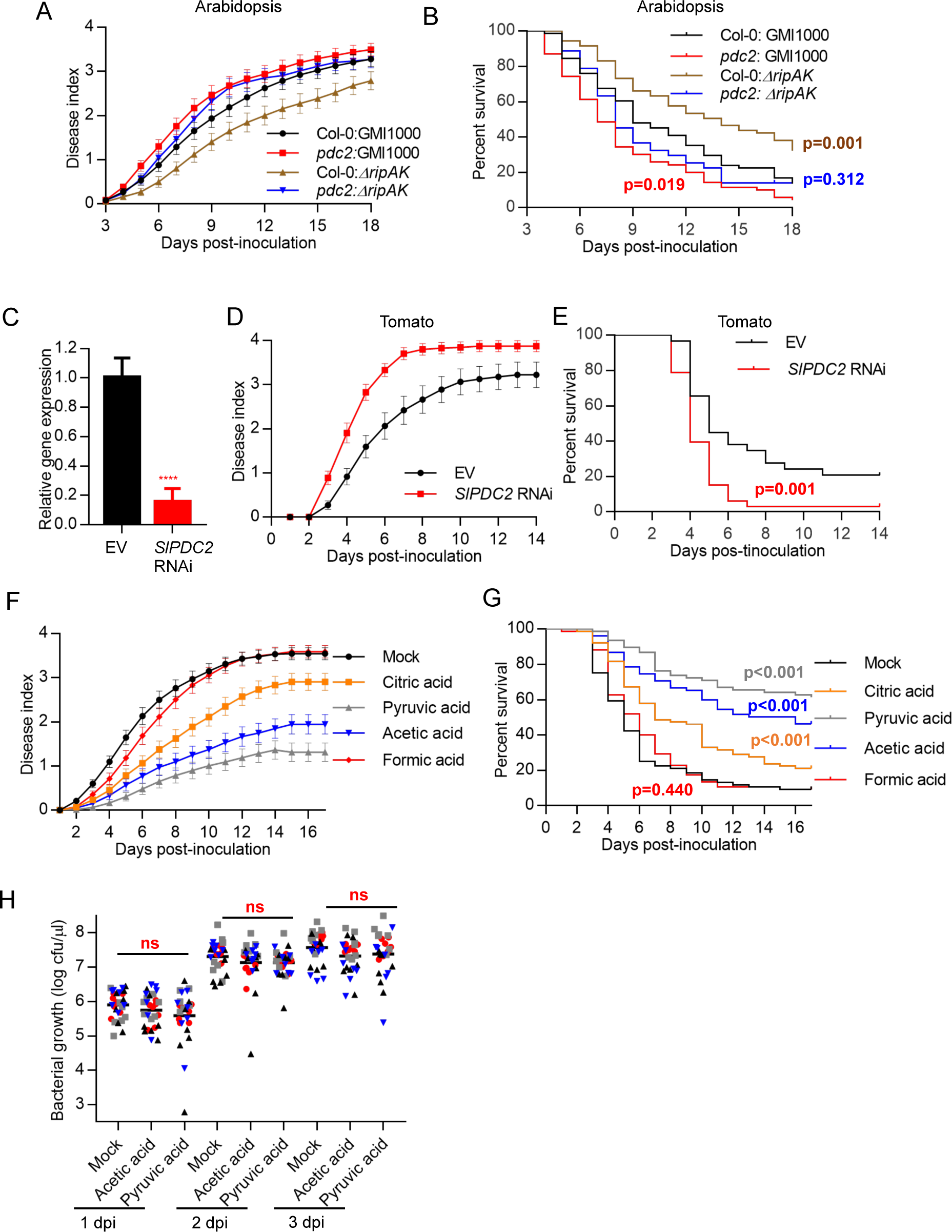
PDCs contribute to plant resistance against *R. solanacearum*. (A and B) The Arabidopsis *pdc2* mutant shows enhanced susceptibility to *R. solanacearum* infection, and rescues the virulence attenuation of the *ΔripAK* mutant. Composite data from 4 independent biological repeats (a representative assay is shown in Figure 3C and 3D). All values were pooled together and represented as disease index (A) or percent survival (B). Disease index values represent mean ± SEM (n=71). (C) Expression of the *SlPDC2* gene in tomato roots expressing the *SlPDC2*-RNAi construct used in the experiments shown in Figure 3E and 3F, determined by qRT-PCR. Values were normalized to the expression of the *SlEF1α-1* gene, and are shown as relative to the expression in roots expressing the empty vector (EV). Values represent mean ± SEM (n=3 samples per genotype), and asterisks represent significant differences according to a Student’s t test (****P<0.0001). (D and E) *SlPDC2* contributes to resistance against *R. solanacearum* infection in tomato. Composite data from 3 independent biological repeats (a representative assay is shown in Figure 3E and 3F). All values were pooled together and represented as disease index (D) or percent survival (E). Disease index values represent mean ± SEM (n=32). Statistical analysis was performed using a Log-rank (Mantel-Cox) test, and the corresponding p value is shown in the graph with the same colour as each curve. (F and G) Soil-drenching inoculation assays in tomato plants upon pre-treatment with a 30 mM solution of the indicated organic acids or water (as mock control). Treatments were performed by placing the pots on a layer of wet towel paper containing the organic acids for 9 days, and then washed and watered normally without treatment for 3 days before inoculation with *R. solanacearum* GMI1000 WT. Composite data from 7 independent biological repeats (a representative assay is shown in Figures 3G and 3H). All values were pooled together and represented as disease index (F) or percent survival (G). Disease index values represent mean ± SEM (n=74). Statistical analysis was performed using a Log-rank (Mantel-Cox) test, and the corresponding p value is shown in the graph with the same colour as each curve. (H) Treatment with pyruvic acid or acetic acid (performed as in F) causes no differences in the growth of *R. solanacearum* GMI1000 upon stem injection. After treatments, 3.5-week old tomato plants were injected with 5 µL of a 10^6^ cfu mL^-1^ and samples were collected 1, 2, and 3 dpi. Four independent biological repeats were performed (n=7 plants per treatment) with similar results. Values from all the replicates are represented in this graph; values with the same colour correspond to the same repeat. ns indicates no significant differences among these treatments according to a Student’s t test (p>0.05).

**Table S1.**
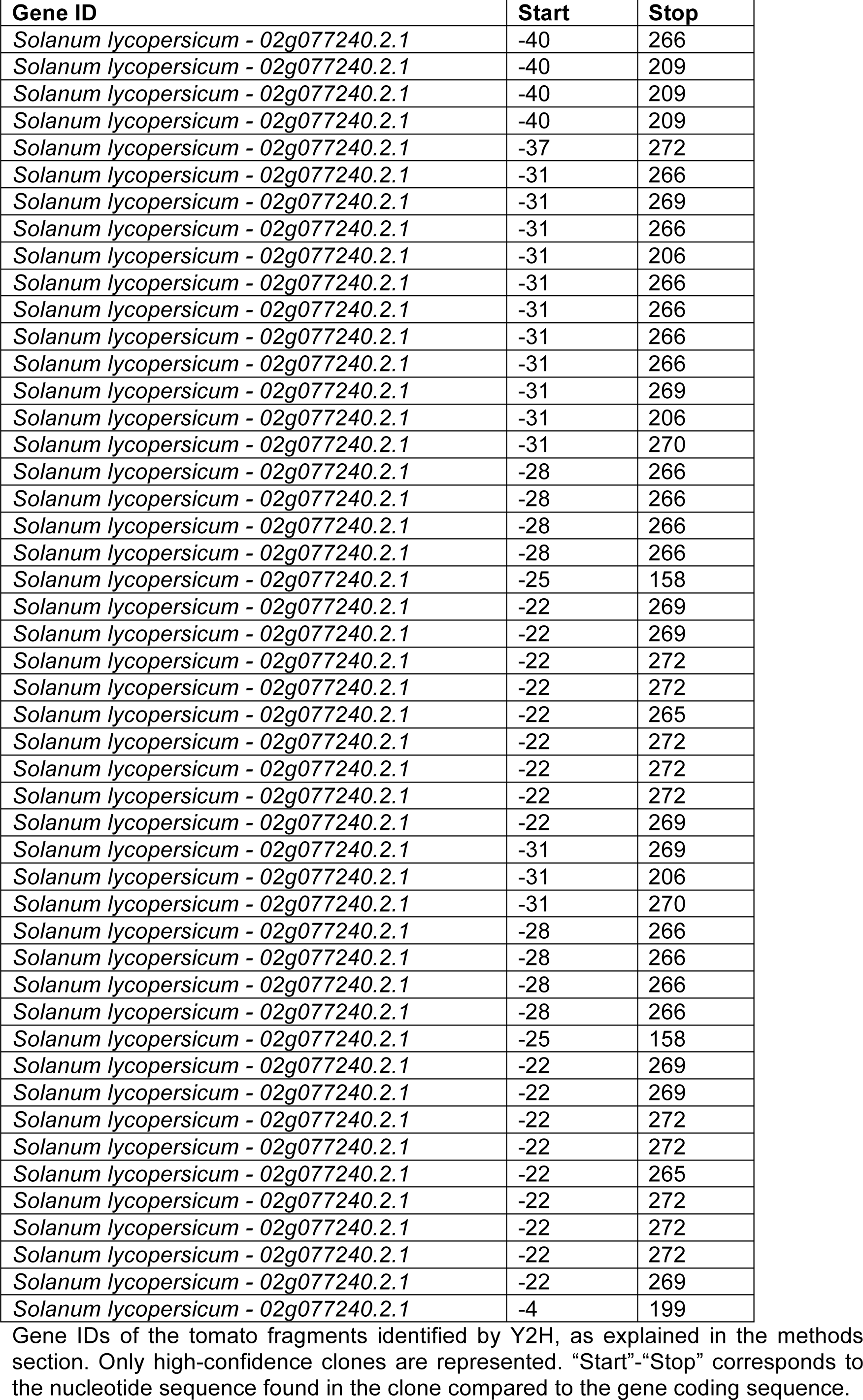
Pyruvate decarboxylase clones identified in the Y2H screen.

**Table S2.**
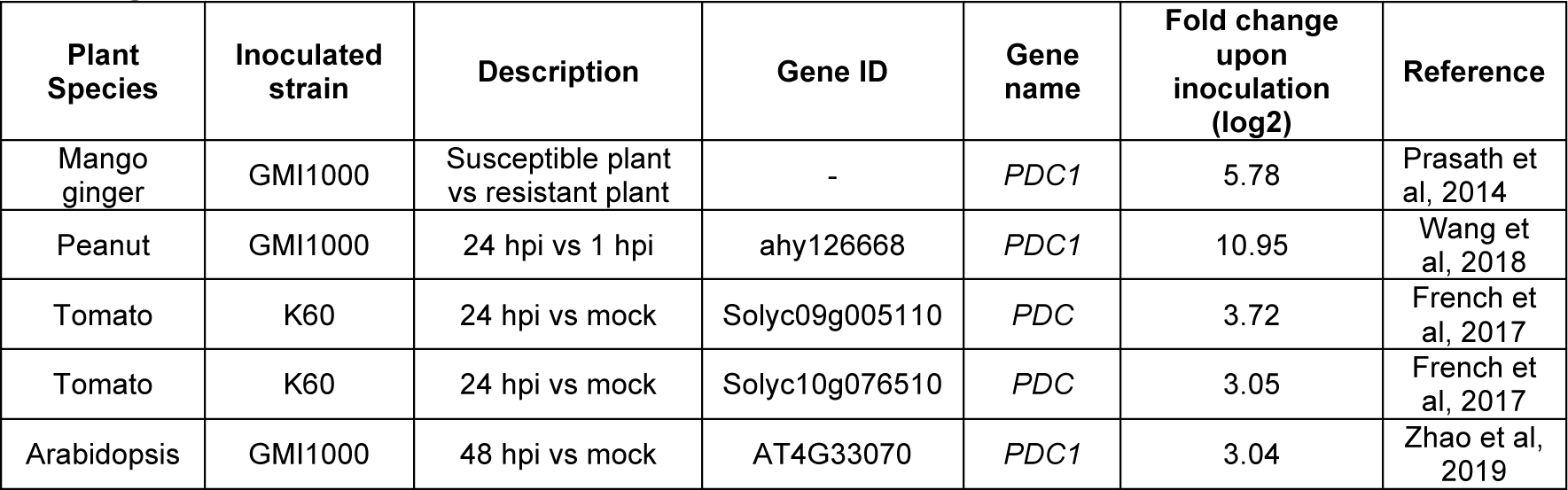
The expression of several PDC orthologs in different plant species is up-regulated upon R. solanacearum inoculation.

**Table S3.**
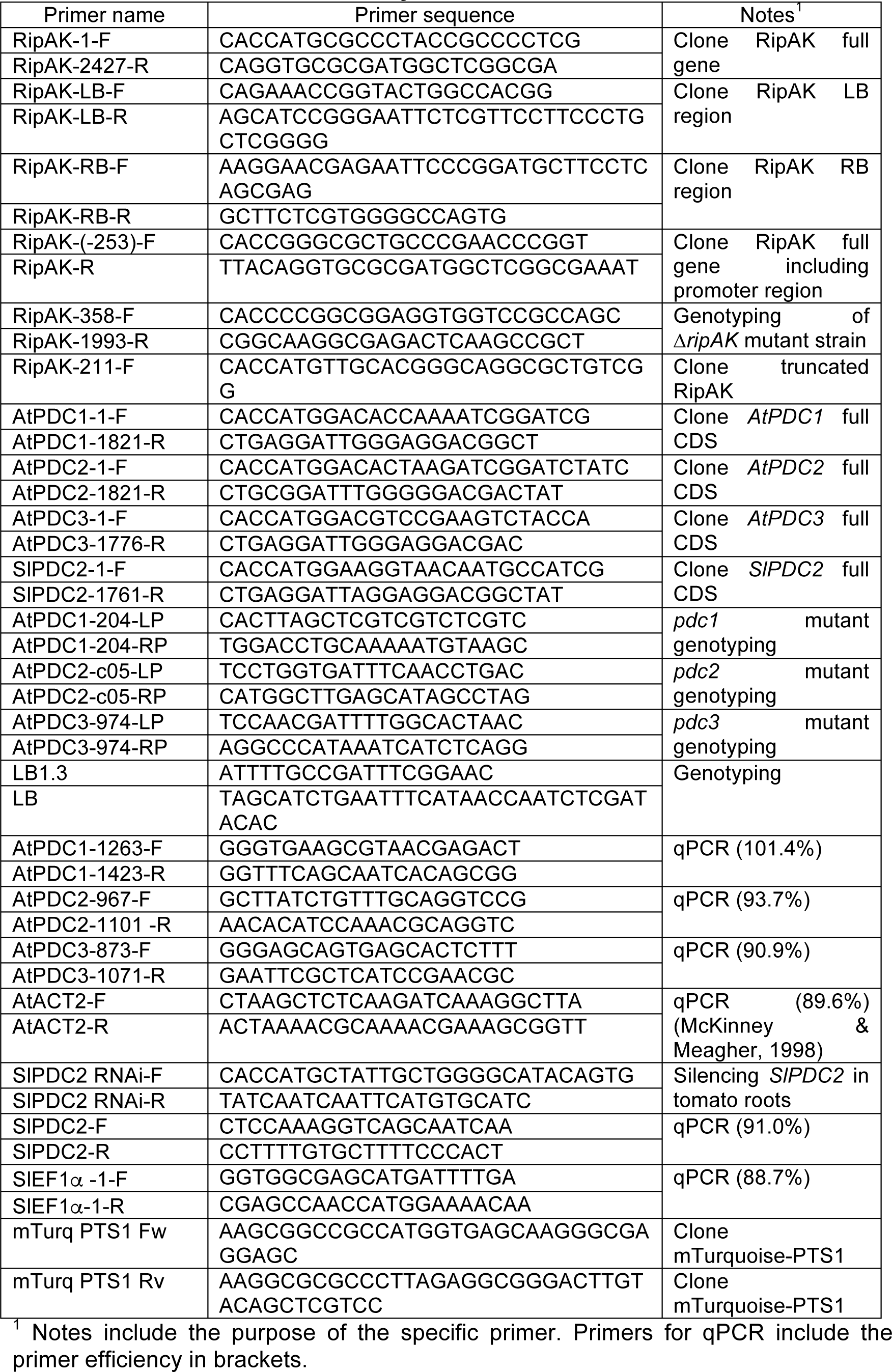
Primers used in this study

